# PRC2 bridges non-adjacent nucleosomes to establish heterochromatin

**DOI:** 10.1101/795260

**Authors:** Rachel Leicher, Eva J. Ge, Xingcheng Lin, Matthew J. Reynolds, Thomas Walz, Bin Zhang, Tom W. Muir, Shixin Liu

**Author notes:** These authors contributed equally to this work (Eva J. Ge, Xingcheng Lin, Matthew J. Reynolds). Correspondence (B.Z.); (S.L.).

## Abstract

Polycomb repressive complex 2 (PRC2) maintains transcriptionally silent heterochromatin by installing and spreading repressive histone methylation marks along nucleosome arrays. Despite extensive research, the mechanism by which PRC2 engages with chromatin remains incompletely understood. Here we employ single-molecule force spectroscopy and molecular dynamics simulations to dissect the interactions of PRC2 with polynucleosome substrates. Our results reveal an unexpectedly diverse repertoire of PRC2 binding configurations on chromatin. Besides interacting with bare DNA, mononucleosomes, and neighboring nucleosome pairs, PRC2 is also found to bridge non-adjacent nucleosomes, an activity associated with chromatin compaction. Furthermore, the distribution and stability of these PRC2–chromatin interaction modes are differentially modulated by accessory PRC2 cofactors, oncogenic histone mutations, and the methylation state of chromatin. Overall, this work provides a paradigm for understanding the physical basis of epigenetic maintenance mediated by Polycomb group proteins.

## INTRODUCTION

Eukaryotic chromatin is modified by a plethora of epigenetic machineries that add or erase specific histone posttranslational modifications, thereby exerting transcriptional control (*1*). Some of these complexes are recruited to and stimulated by their own enzymatic products, which leads to the modification of neighboring nucleosomes and eventually the formation of large-scale chromatin domains. One preeminent example for such a positive feedback mechanism is mediated by the Polycomb repressive complex 2 (PRC2), which catalyzes methylation of the lysine 27 residue on histone H3 (H3K27), resulting in the final enzymatic product, trimethylated H3K27 (H3K27me3), a hallmark of facultative heterochromatin (*2*). PRC2 maintains transcriptional silencing of a large number of genes involved in development and cancer (*3*). The PRC2 core complex comprises four subunits: EED, EZH2, RBBP4, and SUZ12. In the proposed “read-and-write” model, EED recognizes an existing H3K27me3 mark and allosterically activates the methyltransferase EZH2, which in turn modifies a neighboring nucleosome (*4, 5*). This model is supported by a recent cryo-electron microscopy (EM) structure of PRC2 engaging a dinucleosome, in which the EED subunit interacts with an H3K27me3-containing nucleosome and the EZH2 subunit of the same PRC2 complex engages an adjacent unmodified nucleosome (*6*). However, due to the prohibitive conformational heterogeneity of polynucleosome arrays, the binding configuration of PRC2 on its natural chromatin substrate has thus far been refractory to structural interrogation. Moreover, the activity of the PRC2 core complex is influenced by diverse regulatory factors, including accessory subunits, histone modifications, DNA methylation, and RNA [reviewed in (*7, 8*)]. How PRC2 integrates signals from these regulatory factors to achieve chromatin targeting and H3K27me3 spreading remains incompletely understood. To address these knowledge gaps, in this work we developed experimental and computational approaches to probe the binding modes of PRC2 on individual polynucleosome substrates.

## RESULTS

### A single-molecule platform to dissect PRC2–chromatin interactions

We constructed a DNA template harboring twelve repeats of the “601” nucleosome positioning sequence, each separated by 30 base pairs (bp) of linker DNA. After reconstitution with wild-type unmodified histone octamers (Figure S1), individual nucleosome arrays were tethered and mechanically stretched on a dual-trap optical tweezers instrument (Figure 1A). Single-molecule force-extension curves displayed signature sawtooth patterns (Figure 1B) consistent with previous results (*9*), with each abrupt transition signifying the unwrapping of a single nucleosome. The average force at which these transitions occurred is 16.0 ± 0.3 pN (mean ± s.e.m.) and the average number of transitions observed on each array is 12 ± 1. The contour length change (Δ*L*_0_) per transition, which reports the amount of DNA released by force-induced disengagement, is 76 ± 1 bp, in good agreement with the unraveling of the inner DNA wrap around the histone octamer (Figures 1C and S2). The unpeeling of the outer DNA wrap occurs in a gradual fashion at low forces (< 5 pN) under our experimental conditions (*9, 10*).

**Figure 1.**
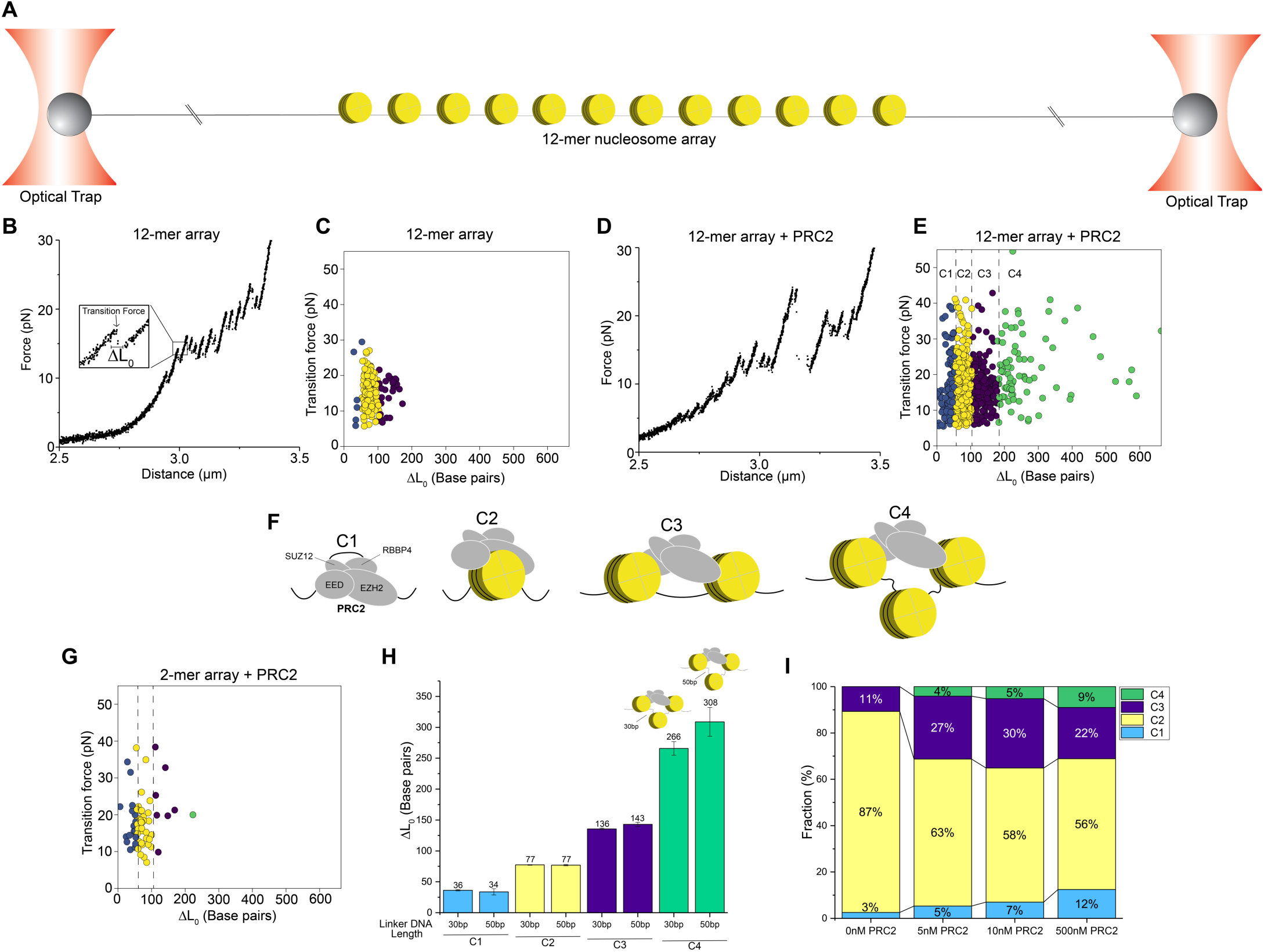
Single-molecule force spectroscopy dissects PRC2 binding modes on a polynucleosome array. (**A**) Schematic of the dual-trap optical tweezers setup. A 12-mer nucleosome array is tethered to a streptavidin-coated bead via a 4-kb biotinylated DNA handle on each side. (**B**) A representative force-extension curve for a 12-mer array and the zoomed-in view of a force-induced transition. Δ*L*_0_ represents the change in contour length after the transition. (**C**) Cluster analysis of all transitions observed in the force-extension curves of individual 12-mer arrays. (**D**) A representative force-extension curve for a 12-mer array incubated with PRC2. (**E**) Cluster analysis of transitions for PRC2-bound 12-mer arrays. (**F**) Cartoon illustrations of the four PRC2-chromatin binding modes representing the four clusters of transitions. C1 shows PRC2 binding to bare DNA; C2 shows PRC2 binding to mononucleosomes; C3 shows PRC2 engaging with two neighboring nucleosomes; C4 shows PRC2 bridging a pair of non-adjacent nucleosomes. (**G**) Cluster analysis of transitions found in PRC2-bound dinucleosome substrates. (**H**) Comparison of the average Δ*L*_0_ for transitions in each cluster between PRC2-bound 12-mer arrays with 30-bp linker DNA versus those with 50-bp linker DNA. (**I**) Cluster distributions for 12-mer arrays incubated with different concentrations of PRC2.

Next, we incubated the 12-mer arrays in a relaxed state (i.e., under zero force) with a saturated concentration of PRC2 core complexes (500 nM) and then subjected them to single-molecule pulling. Interestingly, force-extension trajectories for these assemblies exhibited transitions of diverse sizes (Figure 1D), in contrast to the uniform transitions observed in the absence of PRC2. Using a clustering algorithm based on the Bayesian information criterion (Figure S3), we categorized all transitions into four distinct clusters (Figure 1E). The first cluster (C1) includes transitions smaller than 53 bp, with an average Δ*L*_0_ of 36 ± 2 bp. Considering that PRC2 is known to bind free DNA (*11, 12*), we posited that PRC2 sequesters, perhaps bends, a stretch of free DNA located in the flanking or linker DNA regions of the nucleosome array (Figure 1F, “C1”), and that the C1 transitions are due to force-induced dissociation of this stretch of DNA from PRC2. This interpretation is supported by the force-extension curves for bare DNA substrates incubated with PRC2, which exhibited exclusively these small transitions (Figure S4).

The second cluster (C2) contains transitions between 54 and 107 bp in size with an average Δ*L*_0_ of 77 ± 1 bp (Figure 1E), mimicking transitions in the no-PRC2 condition (Figure 1C) and thus representing the unraveling of mononucleosomes. The transition force was not significantly altered by the presence of PRC2 (16.2 ± 0.3 pN without PRC2 versus 16.6 ± 0.3 pN with PRC2), but was moderately increased by the addition of S-adenosylmethionine (SAM)—the methyl donor for PRC2-catalyzed H3K27 methylation (17.4 ± 0.3 pN with PRC2 and SAM), indicating that the docking of SAM to EZH2 stabilizes its interaction with the nucleosome. While we cannot rule out that some of the C2 transitions may correspond to the unraveling of nucleosomes not bound to PRC2, the saturated amount of PRC2 used in this experiment and the SAM dependence suggest that this cluster of transitions for the most part corresponds to the disassembling of PRC2-bound mononucleosomes (Figure 1F, “C2”).

The third cluster (C3) encompasses transitions ranging from 108 to 180 bp in size (Figure 1E). Besides the fraction that can be attributed to two mononucleosomes coincidentally unraveling at the same time [also seen in the no-PRC2 case (Figure 1C) but occurring with a much lower frequency: 22% with PRC2 versus 11% without PRC2], this cluster also reflects the binding mode in which PRC2 simultaneously engages two neighboring nucleosomes (Figure 1F, “C3”) as seen in the cryo-EM structure (*6*). The rationale is as follows: disruption of the PRC2-dinucleosome complex would release approximately 180 bp of DNA—two outer DNA wraps (∼150 bp) plus 30 bp of linker DNA; alternatively, it would release between 105 and 180 bp of DNA if all or part of the outer wrap from one of the PRC2-interacting nucleosomes is not subjected to PRC2 sequestration, hence already undone at a low force (Figure S5A).

Surprisingly, we also observed a significant population of transitions (C4) with Δ*L*_0_ greater than 180 bp (Figure 1E). These transitions are too large to be explained by any of the interaction modes described above. Rather, they must involve concurrent engagement of PRC2 with two non-adjacent nucleosomes. Specifically, if PRC2 bridges two nucleosomes that are separated by one spacer nucleosome (“Nucl_1-3_” mode), disruption of this linkage would release between 210 and 285 bp of DNA (Figure S5B). If PRC2 bridges a pair of nucleosomes that are separated by two or more spacer nucleosomes (Nucl_1-4_, Nucl_1-5_, etc.), transitions larger than 300 bp would be expected (Figure S5C). Because the great majority of the C4 events (78%) were between 180 and 300 bp in size, we conclude that Nucl_1-3_ is the preferred mode of PRC2-nucleosome engagement in this cluster (Figure 1F, “C4”).

### Validation of the capacity of PRC2 to bridge non-adjacent nucleosome pairs

The interpretation for C4 transitions (i.e., PRC2 bridging two non-neighboring nucleosomes) observed in the force-extension curves of PRC2-bound 12-mer arrays makes a few testable predictions. First, since this mode of PRC2 interaction requires a substrate harboring at least three nucleosomes, it should not be observed on arrays containing only two nucleosomes. We therefore constructed a dinucleosome substrate (Figure S2B) and, satisfyingly, found that C4 was essentially abrogated when pulling on this substrate (Figure 1G), in contrast to the result obtained for the 12-mer substrate (Figure 1E).

Second, the amount of DNA released per C4 transition should be dependent on the linker DNA length. To test this prediction, we constructed a 12-mer nucleosome array with 50-bp linker DNA and conducted single-molecule pulling experiments with this substrate. Our measurements showed that the linker DNA length selectively modulates the size of C4 transitions and that the average Δ*L*_0_ in C4 for the 50-bp-linker array is larger by approximately 40 bp than that for the 30-bp-linker array (Figure 1H). This difference again suggests that the Nucl_1-3_ mode, which releases two linker DNA lengths upon disruption, is the predominant mode for PRC2 engagement in C4.

Third, since the occurrence of C4 transitions strictly relied on the presence of PRC2 (compare Figures 1C and 1E), we expected that the frequency of their occurrence should depend on the amount of chromatin-bound PRC2. We thus conducted pulling experiments with varying concentrations of PRC2 and indeed observed such dependence (Figure 1I). The relative population of C4 transitions, as well as that of C1 transitions representing PRC2–DNA interactions, increased as the PRC2 level was raised from a sub-saturating (5 nM and 10 nM) to a saturating (500 nM) concentration. Taken together, these single-molecule data provide strong evidence that PRC2 can bridge non-adjacent nucleosomes and that this mode of interaction can accommodate different linker DNA lengths.

### *In silico* characterization of PRC2–chromatin interactions

To seek further support for this novel nucleosome-binding activity of PRC2, we performed *in silico* analysis to determine the set of low-energy poses that the PRC2 core complex adopts when in contact with a tetranucleosome substrate. Our analysis combines molecular dynamics simulations to sample tetranucleosome configurations with rigid docking to explore PRC2 binding poses.

We first performed 25 independent molecular dynamics simulations to collect a total of 1,000 representative structures for the tetranucleosome (see *Materials and Methods*). To ensure that both extended and collapsed configurations are included in this structural ensemble, we used harmonic restraints to bias each simulation towards different spatial distances between the 1-3 and 2-4 nucleosome pairs. These simulations were initialized with a tetranucleosome configuration obtained by sequentially extending the dinucleosome cryo-EM structure (*6*) with 30-bp-long linker DNA segments and mononucleosomes. A coarse-grained force field that models protein (*13, 14*) and DNA (*15*) molecules at a single-residue and single-base-pair resolution was used to provide an accurate description of protein–protein and protein–DNA interactions. Prior studies have shown that this level of resolution is sufficient to accurately model the energetics of nucleosomal DNA unwinding (*16, 17*) and inter-nucleosome interactions (*18*). To further improve the computational efficiency, we modeled the core region of each nucleosome as rigid bodies while maintaining the flexibility of the outer layer DNA. As shown in Figure 2A, these simulations covered a wide range of tetranucleosome configurations and allowed us to determine the free energy surface for chromatin folding.

**Figure 2.**
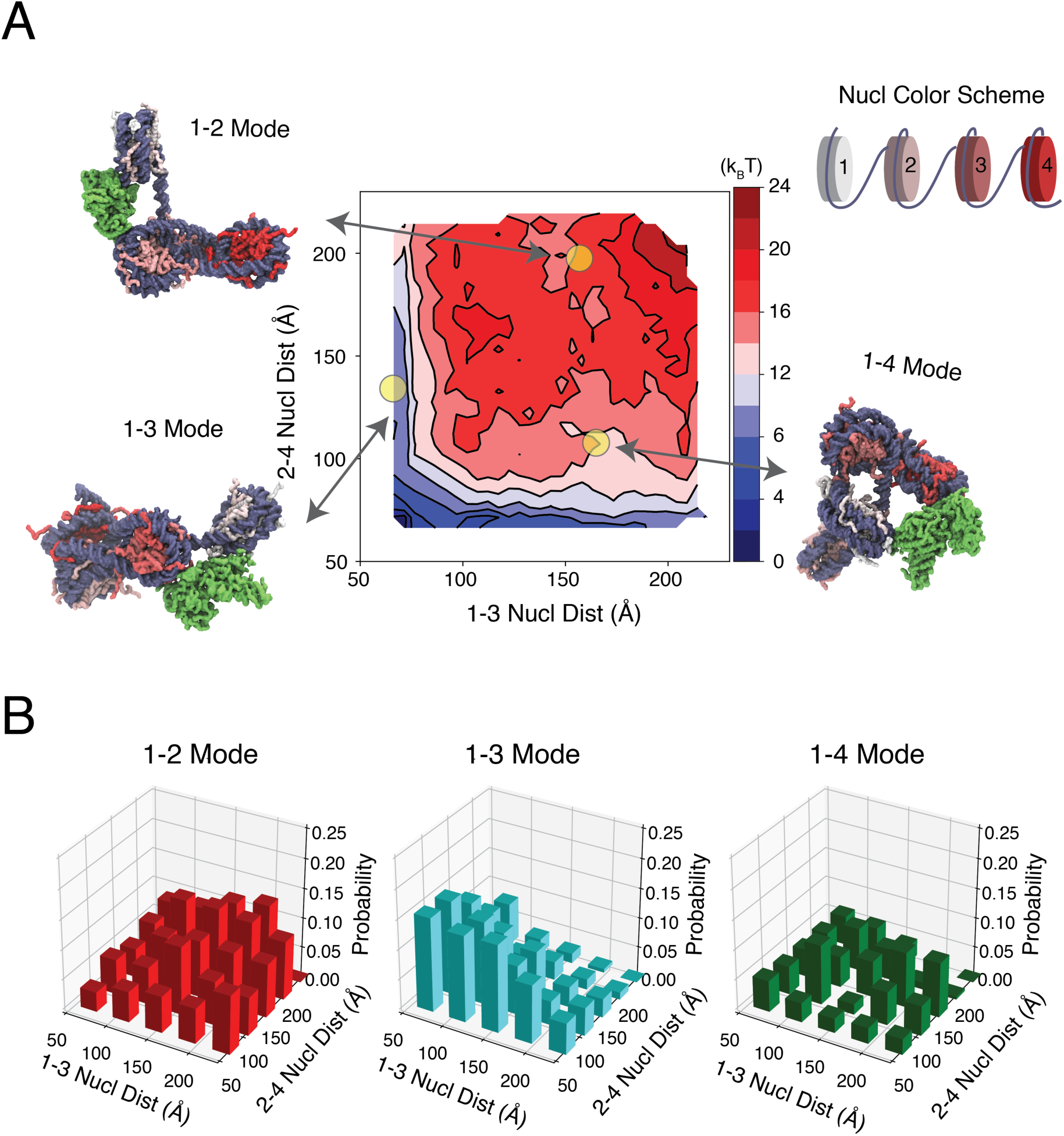
Molecular dynamics simulations reveal the interaction between PRC2 and non-adjacent nucleosomes. (**A**) Thermodynamic stability (free energy) for the large set of tetranucleosome configurations collected for PRC2 docking. Tetranucleosome configurations are grouped by the spatial distances between the 1-3 and 2-4 nucleosome pairs. Example chromatin configurations are shown on the side, with the DNA in purple and the color of histone proteins varying from white to red as the nucleosome index increases. The PRC2 complex is colored in green and shown in its lowest energy pose in the three examples. (**B**) The fraction of PRC2 engaging in various binding modes at different tetranucleosome configurations grouped by the inter-nucleosome distances.

Next, for each one of the tetranucleosome structures, we applied a rigid docking procedure to determine the set of lowest energy PRC2 binding poses. The PRC2 structure, with four core subunits, SUZ12, EZH2, EED, and RBBP4, was constructed using homology modeling (*19*). Close examination of the docking results revealed that, consistent with the published cryo-EM structure (*6*), PRC2 can interact with adjacent nucleosomes using EED and EZH2 subunits (Nucl_1-2_ mode). Repeating the docking procedure using the same system as in the cryo-EM study with a dinucleosome structure led to similar conclusions (Figure S6). There, the computational model indeed succeeded in discovering the experimental configuration as the top-ranking structure. The agreement between simulated and experimental PRC2 binding poses provides strong support for the use of the computational protocol implemented here to study PRC2–chromatin interactions.

Interestingly, at many tetranucleosome configurations, we also found that PRC2 can bridge nucleosome pairs with one (Nucl_1-3_ mode) or two (Nucl_1-4_ mode) spacers using the same two EED and EZH2 subunits. As shown in Figure 2B, for small inter-nucleosome distances, the 1-3 mode is indeed the preferred binding pose. We note that the results in Figure 2B at different inter-nucleosome distances cannot be compared directly with each other since they were obtained by simple counting of the docking results without considering the thermodynamic stability of the PRC2/tetranucleosome complex. With proper thermodynamic re-weighting (see *Materials and Methods*), however, the results can be combined to provide a global estimation of the various PRC2 binding modes across the entire phase space. The relative population of the Nucl_1-3_ mode was estimated to be 18%, in comparison with 4% for the Nucl_1-2_ and 7% for the Nucl_1-4_ mode, respectively. The rest of the population displays the mononucleosome-binding or linker-DNA-binding mode of PRC2. Therefore, the computational modeling, together with the thermodynamic estimation, corroborates the single-molecule data and suggests that bridging non-adjacent nucleosomes is an intrinsic property of the PRC2 core complex.

### Differential regulation of PRC2–chromatin interactions by AEBP2 and JARID2

The activity of the PRC2 core complex is enhanced by the regulatory cofactors AEBP2 and JARID2, which function by either facilitating chromatin targeting or stimulating the catalytic efficiency of the complex (*12, 20–23*). Next, we used the single-molecule experimental platform to assess the effects of these cofactors on the binding configuration of PRC2 on chromatin. Strikingly, the addition of a stoichiometric amount of AEBP2 largely abolished C1 transitions, which correspond to free-DNA sequestration by PRC2 (Figure 3A and 3B). Thus, AEBP2 either reduces DNA binding by PRC2, or alters its binding geometry such that it no longer sequesters or bends DNA (i.e., PRC2 dissociation no longer causes a contour length change). We also found that AEBP2 significantly lowered the transition force averaged over all transitions, which mostly represent PRC2–nucleosome interactions (Figure 3C). These changes indicate that AEBP2 reconfigures the chromatin-binding properties of PRC2, destabilizing its engagement with nucleosomes. Such destabilization may facilitate PRC2 migration on polynucleosomes, which explains AEBP2’s stimulatory effect on the H3K27-methylation activity of PRC2.

**Figure 3.**
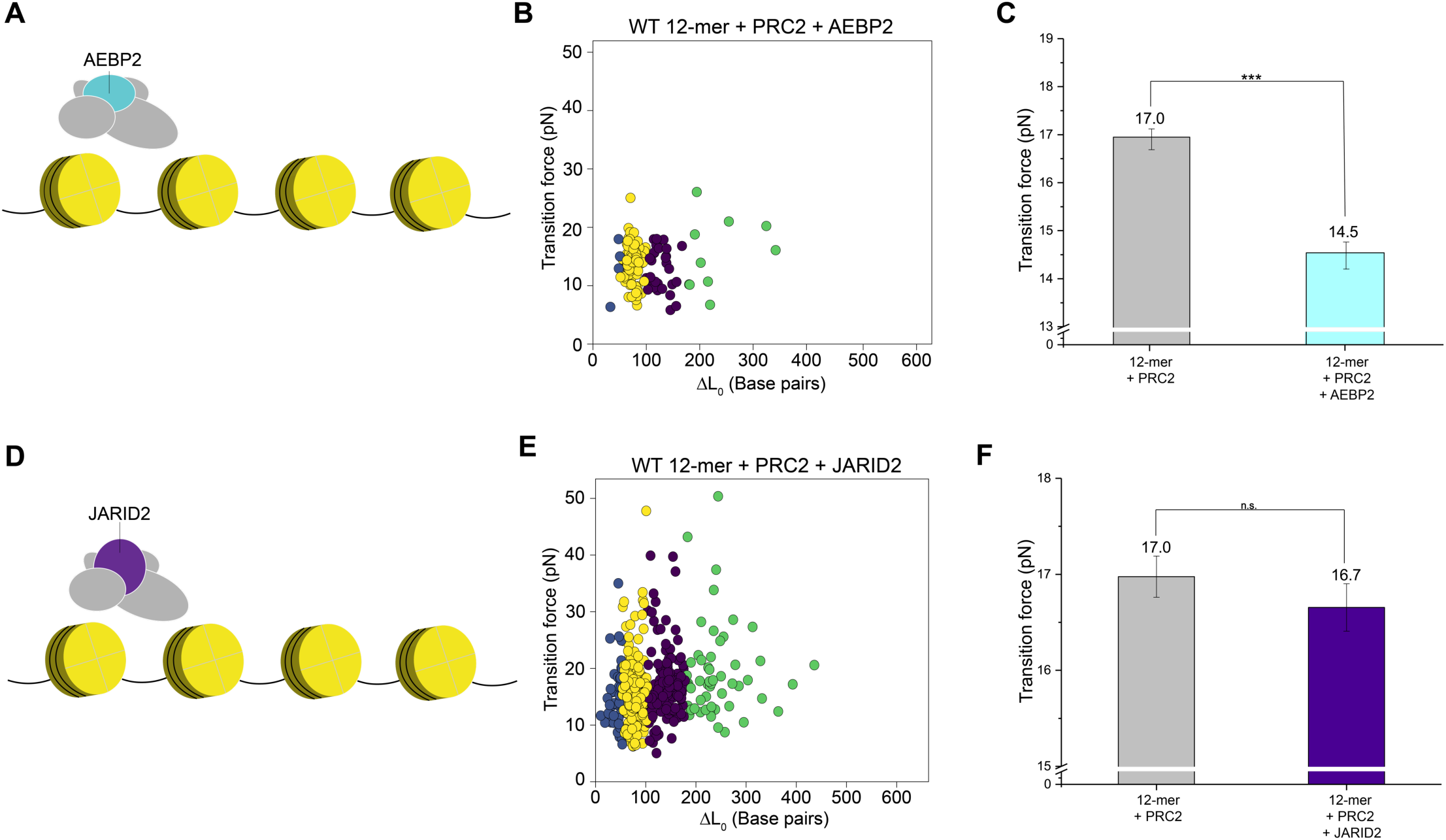
PRC2–chromatin interactions are differentially modulated by AEBP2 and JARID2. (**A**) Cartoon of the PRC2-AEBP2 complex and a nucleosome array. (**B**) Cluster analysis of transitions observed in the force-extension curves for PRC2-AEBP2-bound 12-mer arrays. (**C**) Comparison between the average transition force for 12-mer arrays bound to PRC2 core complexes versus those bound to PRC2-AEBP2 complexes. (**D**) Cartoon of the PRC2-JARID2 complex and a nucleosome array. (**E**) Cluster analysis of transitions observed in the force-extension curves for PRC2-JARID2-bound 12-mer arrays. (**F**) Comparison between the average transition force for 12-mer arrays bound to PRC2 core complexes versus those bound to PRC2-JARID2 complexes.

In contrast, the inclusion of JARID2 did not cause noticeable changes in the distribution of different PRC2–chromatin interaction modes (Figure 3D and 3E), nor did it significantly affect the overall stability of these interactions (Figure 3F). These results are consistent with prior biochemical data (*12*), suggesting that, unlike AEBP2, the stimulatory effect of JARID2 on PRC2’s activity is mainly manifested through allosterically enhancing the catalytic efficiency of EZH2 rather than altering the chromatin-binding behavior of the complex.

### Sensing the histone methylation level of chromatin by PRC2

Previous studies have shown that histone tails and posttranslational modifications therein contribute to the mechanical stability of the nucleosome (*24, 25*). To examine whether preexisting H3K27me3 marks affect the nucleosome stability and/or the nucleosome-binding behavior of PRC2, we constructed 12-mer arrays containing purely H3K27me3-modified nucleosomes, as well as arrays containing an equal mixture of unmodified and H3K27me3-modified nucleosomes (Figure 4A). First of all, single-molecule pulling data showed that the H3K27me3 modification did not significantly affect the stability of nucleosomes themselves (Figure 4B), which is represented here as the strength of contacts between the histone octamer and the inner DNA wrap. However, when pulling on the partially modified arrays that were pre-incubated with PRC2, we observed a significant reduction in the average transition force compared to unmodified arrays (Figure 4C). Remarkably, a similar difference was not detected between unmodified and fully methylated arrays (Figure 4C). This non-monotonic relationship between the methylation level of the substrate and the binding stability of PRC2 suggests that the enzyme can sense the local chromatin modification environment and change its behavior accordingly. PRC2 stably engages with naïve chromatin, which may allow the initial deposition of H3K27me3 marks. The coexistence of activating (modified) and substrate (unmodified) nucleosomes renders PRC2–chromatin interactions more labile, presumably facilitating the spreading of H3K27me3 marks. When the nucleosome array is fully methylated, PRC2 once again becomes stably bound.

**Figure 4.**
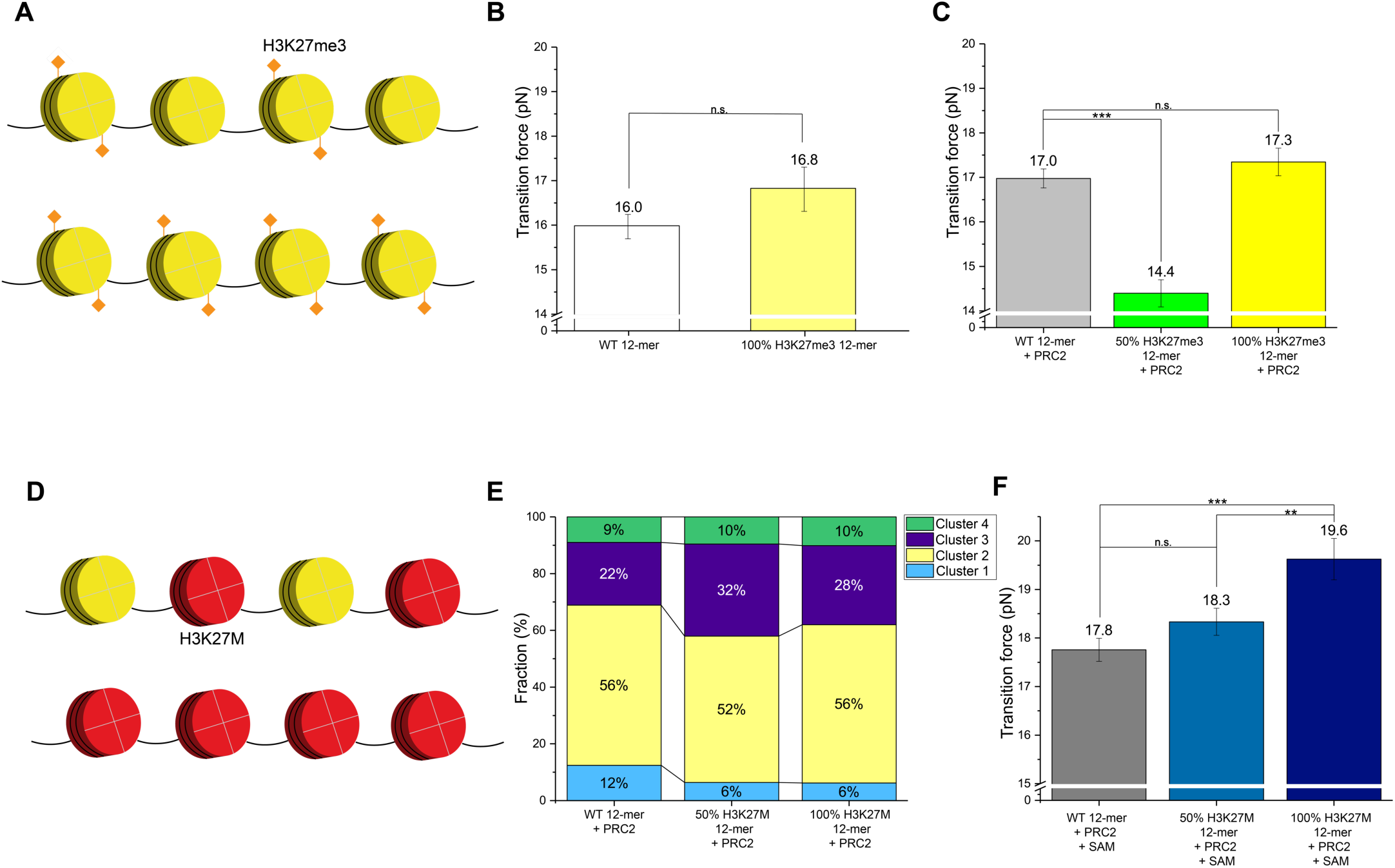
Posttranslational modification and mutation in histone tails affect the behavior of PRC2 on chromatin. (**A**) Cartoon of an array containing 50% (top) or 100% (bottom) H3K27me3-modified nucleosomes. (**B**) The average transition force for wild-type 12-mer arrays and that for 100% H3K27me3-containing 12-mer arrays in the absence of PRC2. (**C**) Comparison of the average transition force for 12-mer arrays containing 0, 50%, or 100% H3K27me3-modified nucleosomes in the presence of PRC2. (**D**) Cartoon of an array containing 50% (top) or 100% (bottom) H3K27M-mutated nucleosomes. (**E**) Cluster distribution for 12-mer arrays containing 0, 50%, or 100% H3K27M nucleosomes incubated with PRC2. (**F**) Comparison of the average transition force for 12-mer arrays containing 0, 50%, or 100% H3K27M nucleosomes in the presence of PRC2 and SAM.

### Effect of an oncohistone mutation on PRC2–chromatin interactions

The lysine-to-methionine substitution of H3K27 (H3K27M) is prevalently found in pediatric brain cancers and is associated with a global decrease of H3K27me3 via inhibiting PRC2 (*26, 27*). To evaluate how this oncogenic mutation impacts PRC2–nucleosome interactions, we compared the force-extension trajectories for PRC2-bound 12-mer arrays that contain 0, 50%, or 100% H3K27M mutant copies of H3 (Figure 4D). A larger overall population of nucleosome-binding clusters (C2, C3 and C4) relative to the DNA-binding cluster (C1) was observed on H3K27M-containing arrays (Figure 4E). Moreover, the mutation stabilizes PRC2 engagement with the chromatin substrate in the presence of SAM, yielding progressively higher transition forces with higher doses of mutant H3 in the array (Figure 4F). This stabilization effect was observed in the mononucleosome binding mode (C2), the adjacent-nucleosome bridging mode (C3), and the non-adjacent-nucleosome bridging mode (C4) (Figure S7). These results hence support the model in which H3K27M traps PRC2 on local nucleosomes via strong contacts, thereby inhibiting the propagation of H3K27me3 modifications (*26, 28, 29*).

### PRC2-mediated compaction of nucleosome arrays

The ability of PRC2 to bridge pairs of non-adjacent nucleosomes not only has implications for the mechanism of H3K27me3 spreading, but it also entails a chromatin-compacting activity of the complex. To test this activity, we examined the architecture of 12-mer nucleosome arrays using negative-stain EM. We found that the 12-mer arrays generally adopted more globular configurations when incubated with PRC2 (Figure 5A and 5B). We then quantified the level of compaction by measuring the average distance between nucleosomes within the same array (Figure S8). This measurement showed that PRC2 condenses 12-mer arrays in a concentration-dependent manner (Figure 5C). We then repeated this EM assay with tetranucleosome substrates and observed the same trend of PRC2-dependent compaction (Figure 5D-5F). These results demonstrate that PRC2 is able to mediate chromatin compaction—a property that can be readily explained by the complex’s distal-nucleosome-bridging activity as revealed by this study. We note that, according to previous work, the chromatin-compacting ability of PRC2 can be further enhanced by replacing the EZH2 subunit with its homolog EZH1 (*30*).

**Figure 5.**
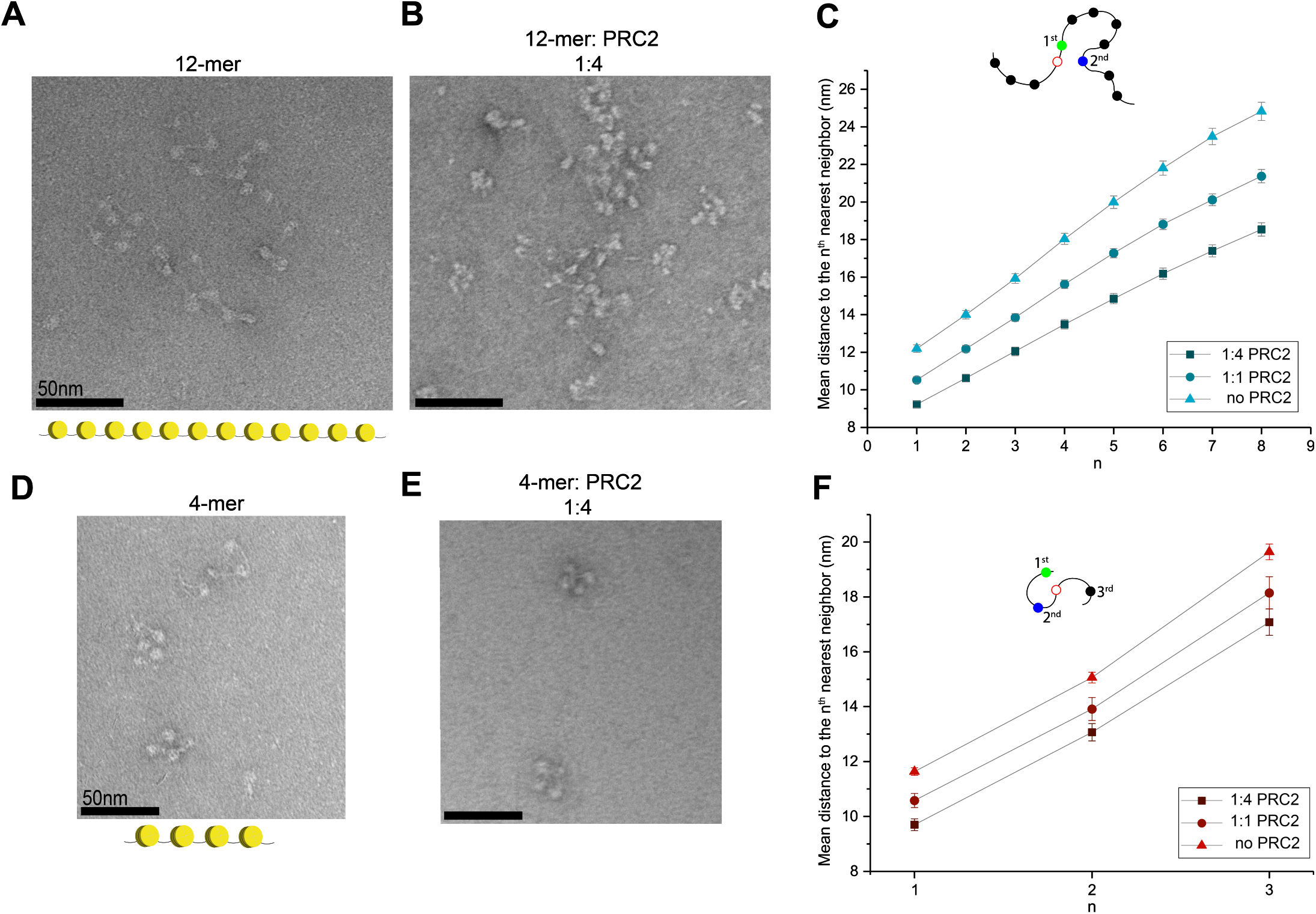
PRC2 mediates the compaction of polynucleosomes. (**A**) A representative negative-stained electron micrograph of 12-mer nucleosome arrays. (**B**) A representative micrograph of 12-mer arrays incubated with PRC2 at a 1:4 ratio (150 nM array and 600 nM PRC2). (**C**) The average distance between a reference nucleosome (open circle) and its *n*^th^ nearest neighbor within the same array measured at different array:PRC2 mixing ratios. A higher PRC2 level correlates to a shorter distance, indicating a higher degree of array compaction. (**D**) A representative electron micrograph of tetranucleosome arrays. (**E**) A representative micrograph of tetranucleosome arrays mixed with PRC2 at a 1:4 ratio. (**F**) The average distance from a given nucleosome (open circle) to its first, second, and third nearest neighbor within the same array measured at different mixing ratios.

## DISCUSSION

By subjecting PRC2–chromatin assemblies to mechanical perturbations, we obtained fresh insights into the chromatin-binding modalities of this important epigenetic machinery (Figure 6). Our data unambiguously demonstrate that PRC2 can bridge non-adjacent nucleosomes and establish the trinucleosome unit as a favored substrate for PRC2 engagement. This conclusion argues against the mechanism in which PRC2 propagates H3K27me3 marks along chromosomes in strictly linear progression. Instead, PRC2 possesses the inherent ability to bypass spacer nucleosomes and perhaps other roadblocks when carrying out the read-and-write function. This feature is required to explain several epigenetic phenomena observed in the cell such as memory and bistability as illustrated by prior theoretical studies (*31, 32*). Such a looping mechanism may be applicable to other chromatin-modifying enzymes, providing flexibility and robustness for their operation (*33*).

**Figure 6.**
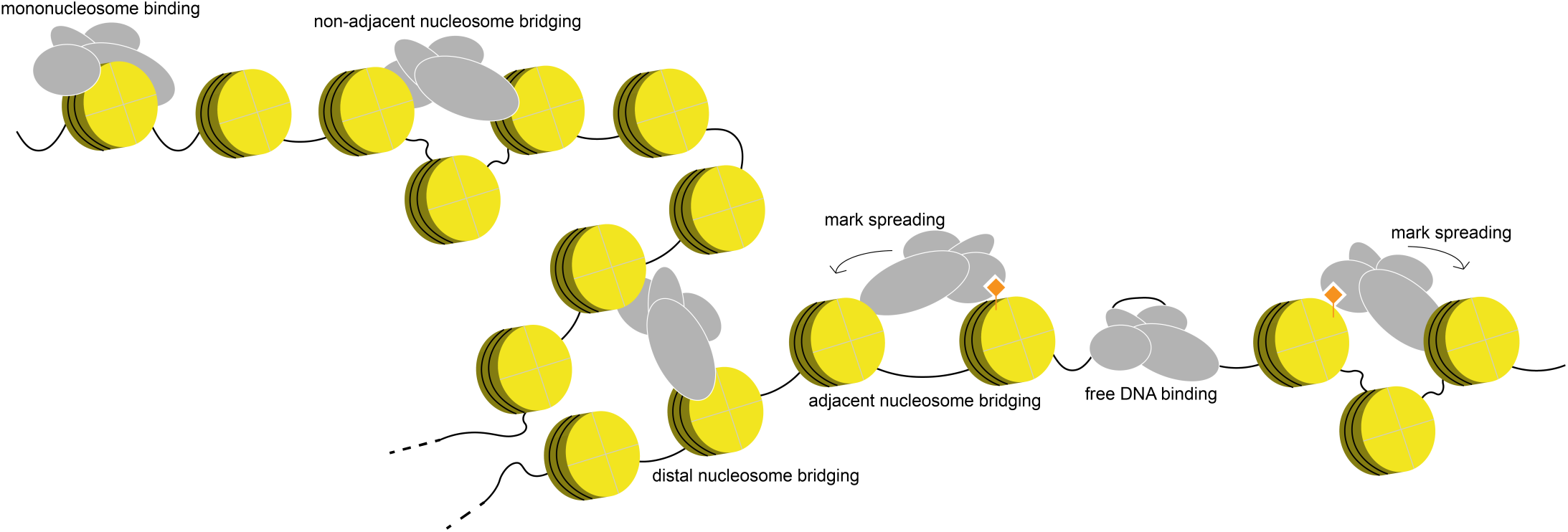
Working model for PRC2-mediated heterochromatin formation. PRC2 interacts with chromatin in multiple modes, resulting in enzymatic modification and non-enzymatic compaction of the chromatin, both of which contribute to transcriptional repression. In particular, the ability of PRC2 to simultaneously engage non-adjacent nucleosomes within the same domain, or perhaps distal nucleosomes from different domains, facilitates efficient spreading of repressive histone marks and mediates local or global chromatin condensation.

Recent studies reported that PRC2–chromatin interactions are transient, allowing the enzyme to readily translocate on the substrate for efficient spreading of repressive marks (*27, 34*). Our data presented here show that PRC2–nucleosome linkages can also withstand high tensions, exhibiting mechanical stability comparable to the nucleosome itself. Although direct evidence is still awaited, it can be envisioned that both stable and transient interactions are at work in order to accomplish de novo installation and self-propagation of H3K27me3 modifications (Figure 6). The strong interactions may be required for PRC2 recruitment to the nucleation sites via either DNA or nucleosome contacts and for the deposition of initial H3K27me3 foci (*34, 35*), presumably with slow kinetics; whereas the weak contacts facilitate efficient spreading by bridging either adjacent or distal nucleosomes. Indeed, we have shown that the mixture of unmodified and modified nucleosomes weakens PRC2 binding to the chromatin, so does the addition of AEBP2. On the other hand, SAM and H3K27M mutation strengthen the PRC2–nucleosome engagement. Our experimental and computational platforms thus provide a framework for delineating the roles of these various regulators—as well as their crosstalk (*36*)—in the balancing act of PRC2.

Finally, the capacity of PRC2 to stably connect non-adjacent nucleosomes also enables it to compact chromatin independent of its enzymatic activity (Figure 6). As such, chemical modification and physical compaction both contribute to Polycomb-mediated heterochromatin formation (*37*). Further experiments to directly visualize the occupancy and movement of PRC2 on chromatin are warranted to answer whether these tasks are accomplished by PRC2 monomers, dimers, or condensates (*38*).

## ACKNOWLEDGMENTS

We thank members of the Liu laboratory for technical assistance and discussions and members of the Walz laboratory for help with electron microscopy. S.L. acknowledges support from the Robertson Foundation, the Quadrivium Foundation, a Monique Weill-Caulier Career Award, a Sinsheimer Scholar Award, a Kimmel Scholar Award, and the National Institutes of Health (DP2HG010510).

## Author contributions

S.L. oversaw the project. R.L. prepared nucleosome and DNA samples, performed the single-molecule experiments, as well as the EM experiments under the supervision of T.W.. E.J.G. and T.W.M. provided the PRC2 samples. M.J.R. developed analysis software and performed EM data processing. X.L. and B.Z. performed computational modeling. R.L. and S.L. wrote the manuscript with input from all authors. Correspondence regarding computational modeling should be addressed to B.Z. All other requests regarding data and materials should be addressed to S.L.

## MATERIALS AND METHODS

### Sample preparation

#### Expression and purification of PRC2 complexes

All PRC2 proteins used in this study were prepared in Sf9 cells using a baculovirus system. Flag-tagged EZH2, and His-tagged EED, RBBP4, and SUZ12 were cloned into a pACEBac1 vector using the Multibac system (*39*). His-AEBP2 and Flag-JARID2 were prepared in their own individual pACEBac1 vectors. Plasmids were used to generate bacmids according to manufacturer protocol (Multibac, Geneva Biotech). 2.5 µg of bacmid was transfected into 1.0 × 10^6^ attached Sf9 cells in a 6-well plate. Following transfection, cells were overlaid with 2 mL of fresh medium (Sf-900III SFM, Thermo Fisher Scientific) and incubated at 27 °C for 96 h in the dark. The supernatant was collected, filtered through 0.22 µm, and supplemented with 2% v/v fetal bovine serum (FBS) to produce the P1 virus. P2 virus was generated by infection of 10 mL of Sf9 cells (1.5 × 10^6^ cells/mL) with 1 mL of P1 virus solution. Cells were grown at 27 °C in suspension culture until they reached <50% viability as monitored by trypan blue staining. Culture supernatant was collected, filtered, and supplemented with 2% v/v FBS. To generate the P3 virus, 300 µL of P2 virus solution was added to 50 mL of Sf9 cells (1.5 × 10^6^ cells/mL). Cells were grown at 27 °C in suspension culture until they reached <50% viability. Culture supernatant was collected, filtered, and supplemented with 2% v/v FBS.

For protein expression, a 1:100 dilution of P3 virus was added to Sf9 cells at 2.0 × 10^6^ cells/mL density. After 48 h of incubation at 28 °C in the dark, cells were harvested by centrifugation and lysed by a Dounce homogenizer (Wheaton) in HEGN600 buffer (25 mM HEPES pH 7.0, 600 mM NaCl, 1 mM EDTA, 10% v/v glycerol, 0.02% NP-40). Soluble extracts were incubated with anti-Flag M2 affinity gel (100 µL resin per 100 mL cell culture) in HEGN350 (25 mM HEPES pH 7.0, 350 mM NaCl, 1 mM EDTA, 10% glycerol, 0.02% NP-40) for 2 h at 4°C. Bound proteins were eluted with HEGN350 containing 0.25 mg/mL Flag peptide (3 × 20 min at 4 °C). Eluted proteins were pooled, spin concentrated in a Vivaspin centrifugal concentrator (MWCO 30,000, Viva Products), and purified by size-exclusion chromatography on a Superose 6 column (GE Healthcare). Fractions containing monomeric PRC2 (as analyzed by SDS-PAGE) were pooled, flash frozen with liquid N_2_, and stored at −80 °C. Prior to use, protein concentration was quantified by A_280_ (PRC2 core complex extinction coefficient = 296,910).

#### Expression and purification of histone proteins

The human core histones (H2A, H2B, H3, and H4) were expressed in BL21 (DE3) cells. The cells were lysed and inclusion bodies were harvested through rounds of sonication and centrifugation (*40*). Histones were extracted from inclusion bodies with DMSO and purified through ion-exchange chromatography. Octamers were reconstituted using salt dialysis and size-exclusion chromatography (Superdex 200 10/300) as described previously (*40*). H3K27me3-modified histone octamers were purchased from EpiCypher. K27M mutant H3 histones were cloned using site-directed mutagenesis.

#### Preparation of DNA templates

DNA templates were digested from a plasmid containing 12 repeats of 601 nucleosome positioning sequences (*41*) in a pET28b backbone, and then ligated to DNA handles containing two biotins on each end.

#### Reconstitution of nucleosome arrays

Nucleosome arrays were formed on the DNA template described above through salt-gradient dialysis from 1.4 M KCl to 10 mM KCl in Slide-a-Lyzer MINI Dialysis units (7,000 MWCO) using a peristaltic pump set at a 1 mL/min flow rate for 6 h. Mouse mammary tumor virus (MMTV) DNA was added during dialysis to prevent octamer overloading.

### Single-molecule experiments

#### Data acquisition

Single-molecule experiments were performed at room temperature on a LUMICKS C-Trap instrument equipped with dual-trap optical tweezers (*42*). A computer-controlled stage enabled rapid movement of the optical traps within a five-channel flow cell. DNA tethers were formed between two 3-µm streptavidin-coated polystyrene beads (Spherotech) held in two laser traps. The tethers were subjected to mechanical pulling by moving one trap relative to the other at a constant velocity (0.1 µm/s), generating force-extension curves. Data were collected at 50 kHz. All experiments were performed with Imaging Buffer (10 mM Tris-HCl pH 8.0, 0.1 mM EDTA, 200 mM KCl, 0.5 mM MgCl_2_, 0.1% Tween, and 1 mM DTT). Unless noted otherwise, saturating concentrations (500 nM) of PRC2, PRC2-AEBP2, or PRC2-JARID2 were used in these experiments.

#### Force-extension analysis

Force-extension traces were processed using a custom Force-Extension Analyzer software suite. Each trace was separated into segments of constant contour length (*L*_0_) based on disruption peaks. Disruption peaks were identified by applying a Butterworth low-pass filter to the trace and assigning transitions when the force decreased by a user-specified threshold, typically 0.2 pN. The unfiltered data for each segment were then fit with the modified worm-like-chain (WLC) model: 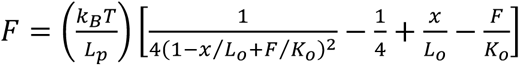 (*43*). In order to prevent over-fitting and to allow for direct comparison of the array’s changes in *L*_0_, the persistence length (*L*_p_) and elastic modulus (*K*_0_) were determined for each nucleosome array based on the first and last segments, respectively. *L*_p_ was determined by fitting the Marko-Siggia WLC model to the first segment in the low force regime (*44*). Keeping this *L*_p_ fixed, the array’s *K*_0_ was determined by fitting the last segment, in the high force regime, to the modified WLC model. Each segment was subsequently fit to the modified WLC model to determine its *L*_0_, maintaining *L*_p_ and *K*_0_ constant for a given trace. Relevant measurements (number of segments, transition force, contour length change Δ*L*_0_) for each trace were exported. The code and tutorial of the Force-Extension Analyzer software is available at https://github.com/alushinlab/ForceExtensionAnalyzer.

#### Cluster assignment

In order to identify the groups composing all observed transitions, the distribution of changes in contour length was modeled using a Gaussian mixture model. Putative models were generated with varying numbers of components, and each model was assessed using the Bayesian information criterion (BIC) score (Figure S3). Models with fewer than four components had substantially worse BIC scores than those with four or more components. The models with four or five components had similar BIC scores and virtually the same cluster boundaries, with the only effective difference being that the cluster of large transitions would be further subdivided in the five-component model. Therefore a four-component model was used in subsequent analyses. Individual transitions were assigned to one of the four clusters based on this probabilistic model. Modeling was implemented in Python using the Scikit-learn library (*45*).

### Computer modeling and simulations

#### Structural modeling for PRC2 and tetranucleosome

We built a structural model for the full length human PRC2 protein using homology modeling (*19*). Only the four core subunits, SUZ12, EZH2, EED, and RBBP4 were included in the structure. Three partially solved EM/X-ray structures of PRC2: PDBID 6C23 (*46*), 5WAI (*47*) and 5HYN (*29*) were used as templates for structural modeling. Missing residues that cannot be found in any of the PDB structures were built as random loops.

An initial configuration for the tetranucleosome was obtained by sequentially extending the dinucleosome cryo-EM structure (*6*). A 30-bp-long linker DNA (sequence: TATGACAGTGCATCACGGGGTGAGATCGCT) was used to connect the 2^nd^/3^rd^ and the 3^rd^/4^th^ nucleosomes. The linker DNA segments were constructed in the perfect helical form using 3DNA tool kit (*48*), and their orientations were dictated by the exiting nucleosomal DNA configuration to minimize bending and twisting. Configurations for the 3^rd^ and 4^th^ nucleosomes were taken from the PDB structure (ID: 3LZ1) and we replaced the protein coordinates with those from 1KX5 to model disordered histone tails. In the final construct, all nucleosomes share the same 601 sequence. We note that our results and conclusions are independent of this initial configuration because of the use of extensive simulations for equilibration.

#### Coarse-grained simulation of tetranucleosomes

We combined the 3SPN.2C DNA model (*15*) and the structure-based C-α model (*13, 49*) to create a coarse-grained force field for accurate and efficient modeling of protein–DNA interactions. We represent each DNA base with three beads and every α-carbon with one bead. Prior studies have shown that this level of resolution is sufficient to accurately model the energetics of nucleosomal DNA unwinding (*16, 17*) and inter-nucleosome interaction (*18*).

The energy function of the system includes contributions from intra-DNA, intra-protein, inter-protein and protein-DNA interactions. Parameters from 3SPN.2C were directly applied to model intra-DNA interactions for the tetranucleosome sequence studied here. When simulating proteins with the structure-based model, we treated each histone octamer as a single unit. Intra-protein interactions therefore refer to all interactions within an octamer, while inter-protein interactions correspond to those between octamers. To ensure the stability of the histone octamer during simulation, we included a list of native contacts for intra-protein interactions. These contacts were generated from the PDB structure (ID: 1KX5) using the Shadow contact map (*50*). Two residues were considered in contact if their minimal atomic distance is 6 Å or less, regardless of whether they are from the same protein chain or not. We scaled the energy of the structure-based model by a factor of 3.75 to 0.9 kcal/mol to keep the protein complex from unfolding at a temperature of 300 K. Detailed expressions of the energy function for the protein and DNA models can be found in Ref. (*14, 15*). Electrostatic interactions modeled at the Debye-Hückel level were included between charged beads, including DNA phosphates, Lys, Arg, Glu, and Asp residues. A salt concentration of 150 mM was used for the screening effect. In addition, a weak, nonspecific Lennard-Jones potential was applied between all protein–DNA beads. Detailed expressions for these potentials can be found in Ref. (*51*).

To further improve the computational efficiency, we modeled the core region of each nucleosome as rigid bodies. This region includes the folded segments of the histone octamer and the central 107 bp of DNA. The rigid units consist of six degrees of freedom for translation and rotation and all atoms within the same unit move concurrently. Our setup maintains the flexibility of the linker DNA, part of the outer-wrap nucleosomal DNA, and disordered histone tails. Since the inner DNA wrap is known to bind tightly to the well-folded histone core in resting nucleosomes under no stress, we anticipate the rigid-body treatment to be a good approximation.

The most stable tetranucleosome configuration is expected to be collapsed under a salt concentration of 150 mM. To explore more expanded configurations in which the tetranucleosome might bind more favorably with PRC2, we carried out 25 independent simulations with the presence of harmonic biases. These biases were implemented to restrain the distances between the non-adjacent nucleosomes (1-3 and 2-4 nucleosomes) at specified values and adopt the following expression:

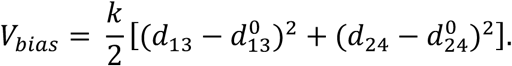

*d*_13_ and *d*_24_ stand for the distance between the 1-3 and 2-4 non-neighboring nucleosomes.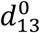 and 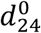 are the corresponding target values and range from 80 to 200 Å with an increment of 30 Å. *k* = 0.01 kcal/mol.

Molecular dynamics simulations were carried out with a time step of 5 fs using the LAMMPS software package to explore tetranucleosome configurations. The Nosé-Hoover thermostat was applied to maintain the simulations at a temperature of 300 K. Periodic boundary condition was enforced with a box size of 2,000 Å. A total of at least 7.5 million steps were carried out for each simulation and we saved the configurations along the trajectory at every 5,000 steps. The first 3 million steps were discarded as equilibration periods.

#### Rigid-body docking for PRC2–tetranucleosome binding

In principle, one can study PRC2 binding with the tetranucleosome via molecular dynamics simulations. However, the slow timescale associated with diffusion and the rugged energy landscape make an exhaustive exploration of different binding modes challenging. To more efficiently study the binding, we, therefore, applied a rigid docking procedure as detailed below.

First, to account for the conformational flexibility of the tetranucleosome, we selected 1,000 structures from biased simulations introduced in the last section. The structures were chosen based on a K-means clustering over all the simulated configurations to include both collapsed and extended configurations. Each configuration was represented with the six inter-nucleosome distances for clustering.

For each one of the 1,000 structures, we then determined the set of most stable PRC2 binding configurations by evaluating the energy of a large set of structures. Specifically, we selected 577 PRC2 orientations from a uniform sampling of the three Euler angles. For each orientation, we then searched for every possible position on a grid of size ∼800×800×800 Å^3^ with a spacing of 2 Å using discrete Fourier transform. Interaction energy between PRC2 and tetranucleosome was evaluated using the same force field introduced in the last section. To avoid steric clashes, we did not use the disordered regions of PRC2 (SUZ12: residue ID 1-78, 150-153, 168-181, 210, 224-227, 254-294, 323-350, 364-422, 549-560, 686-739; EZH2: residue ID 1-9, 183-210, 217-219, 250-256, 346-421, 480-513, 741-746; EED: 1-76; RBBP4: 1-2, 94-104, 413-425) for energy evaluation.

#### Population estimation of PRC2 binding modes

From the docking simulations performed in the last section, we can estimate the fraction of different PRC2 binding modes at a given tetranucleosome configuration. These data, however, cannot be combined straightforwardly to estimate an overall probability for various binding modes as the tetranucleosome configurations were collected from biased simulations. Proper thermodynamic re-weighting must be carried out before averaging as detailed below (*52*).

The probability of PRC2 binding in between adjacent nucleosomes (Nucl_1-2_ mode) at distances 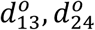 for 1-3 and 2-4 nucleosomes can be defined as:

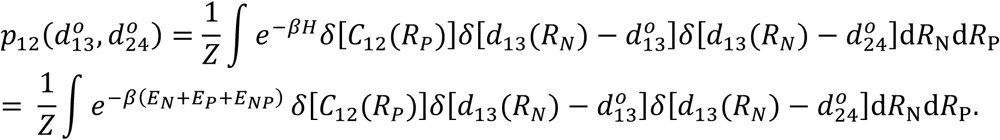

where *R*_N_ and *R*_P_ correspond to the nucleosome and PRC2 degrees of freedom and *z =* ∫*e*^*–βH*^d*R*_N_d*R*_p_ is the partition function. *H* = *E*_N_ + *E*_P_ + *E*_NP_ is the potential energy, with *E*_N_, *E*_P_, and *E*_NP_ being the internal energy of the tetranucleosome, the internal energy of PRC2, and the binding energy between PRC2 and nucleosomes, respectively. δ[*C*_12_(*R*_P_)] represents all the configurations in which PRC2 binds simultaneously with the first and second nucleosomes.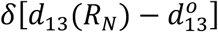 and 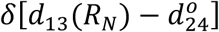 selects out tetranucleosome configurations with 1-3 nucleosome distance at 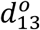 or 2-4 nucleosome distance at 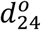, respectively.

The above definition, though formally exact, is difficult to calculate in practice, as it requires simulations in the presence of both PRC2 and tetranucleosomes. To make progress, we invoke the mean field approximation and replace *E*_*NP*_ with the average value at the given distances 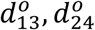 for the Nucl_1-2_ mode, 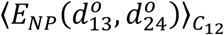. The probability can now be simplified as

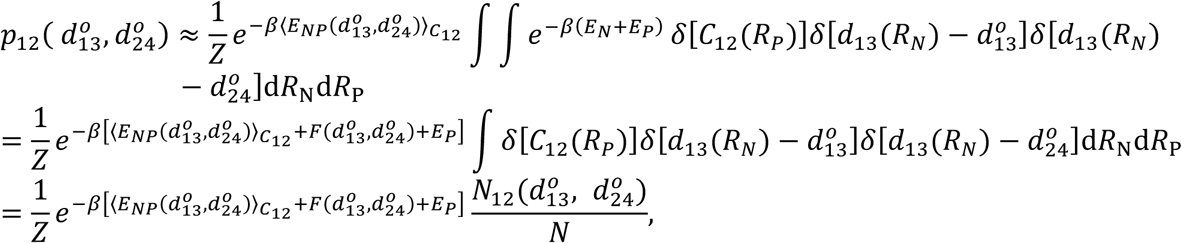

where 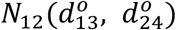 is the number of docked PRC2 structures engaging in Nucl_1-2_ mode at distances 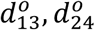, while *N* is the total number of docked PRC2 configurations. 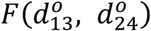 is the free energy of tetranucleosomes with different 1-3, 2-4 nucleosome distances, and was calculated with the weighted histogram analysis method (WHAM) from the biased simulations. To obtain the final population estimation independent of inter-nucleosome distances, we integrate over 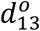 and 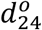

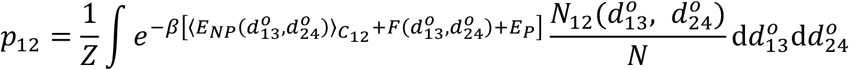

Population of Nucl_1-3_ and Nucl_1-4_ binding modes can be similarly defined as:

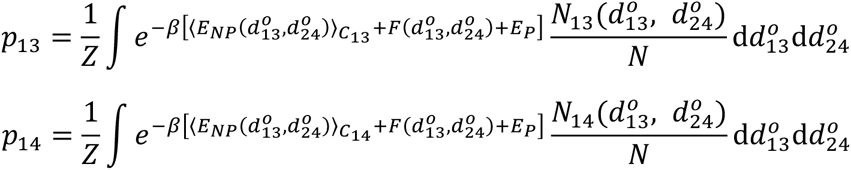

To determine the numerical values of the various populations, the integration was converted into a summation over the 1,000 tetranucleosome structures. For each structure, we used the top 1,000 lowest PRC2 binding configurations determined from docking to estimate 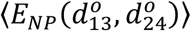 and 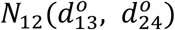. When determining the binding modes, the minimum atomic distance between PRC2 and each of the four nucleosomes was calculated. A cutoff distance of 0.8 nm (approximately one Debye length) was used to determine whether the PRC2 and the corresponding nucleosome was in contact.

### EM analysis

#### Negative-stain electron microscopy

12-mer and 4-mer nucleosome arrays were formed using salt dialysis as described above. Arrays were purified by MgCl_2_ precipitation (4 mM final MgCl_2_ concentration, centrifugation at 15,000 rpm for 10 min at 4 °C). PRC2 and nucleosome arrays were dialyzed into EM buffer (50 mM HEPES pH 7.9, 50 mM KCl, 1 mM TCEP). Samples were adsorbed to glow-discharged carbon-coated copper grids and stained with uranyl formate. Imaging was performed on a CM10 electron microscope at a nominal magnification of 52,000×.

#### Image analysis of electron micrographs

The coordinates of nucleosomes in each micrograph were manually picked using the e2boxer tool in EMAN2 (*53*). These coordinates were assigned to nucleosome arrays by detecting whether they existed within masks encompassing the arrays. These masks were generated for each nucleosome array as follows: negative-stain micrographs were down-sampled after binning over 4×4 pixels to speed up computations. Uneven illumination was corrected by calculating an adaptive, local threshold and subtracting it from the down-sampled micrograph. These images were then entropy-filtered, followed by binarization at a fixed threshold per micrograph. Masks were slightly dilated and returned to the original scale. Picked coordinates within a mask were considered to be part of the same nucleosome array. See Figure S8 for an example micrograph.

### Statistical Analysis

Errors reported in this study represent the standard error of the mean. *P*-values were determined from two-tailed two-sample *t*-tests (n.s., not significant; *, *P* < 0.05; **, *P* < 0.01; ***, *P* < 0.001).

---

**Figure S1.**
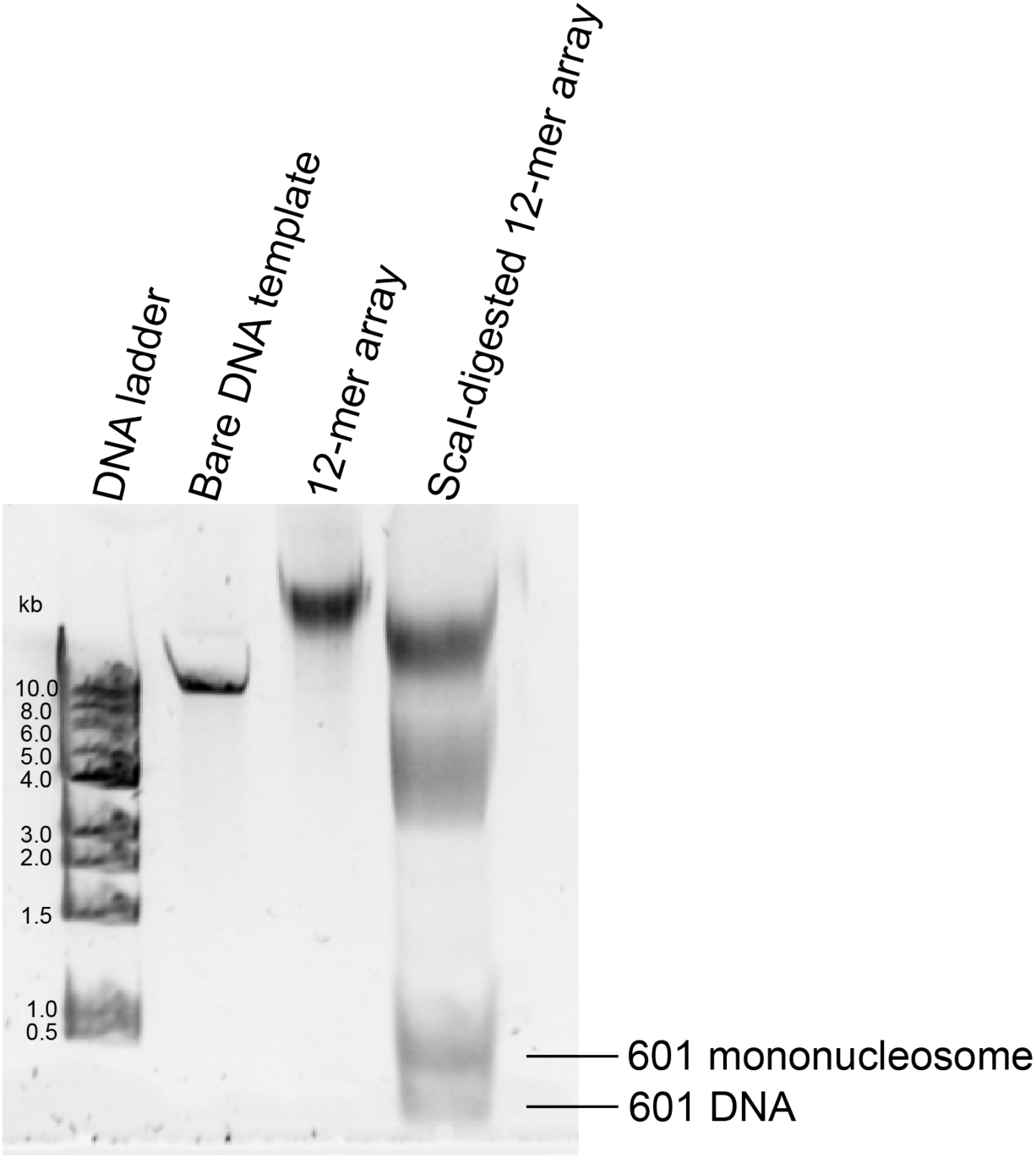
Formation and analysis of 12-mer nucleosome arrays. APAGE gel shows the substrates used for optical-tweezers experiments. Lane 1: DNA ladder; Lane 2: 10-kb bare DNA template containing twelve “601” nucleosome positioning sequences, each separated by a 30-bp linker DNA with a ScaI restriction site; Lane 3: the same DNA template loaded with histone octamers to generate the 12-mer nucleosome array; Lane 4: ScaI-digested 12-mer array.

**Figure S2.**
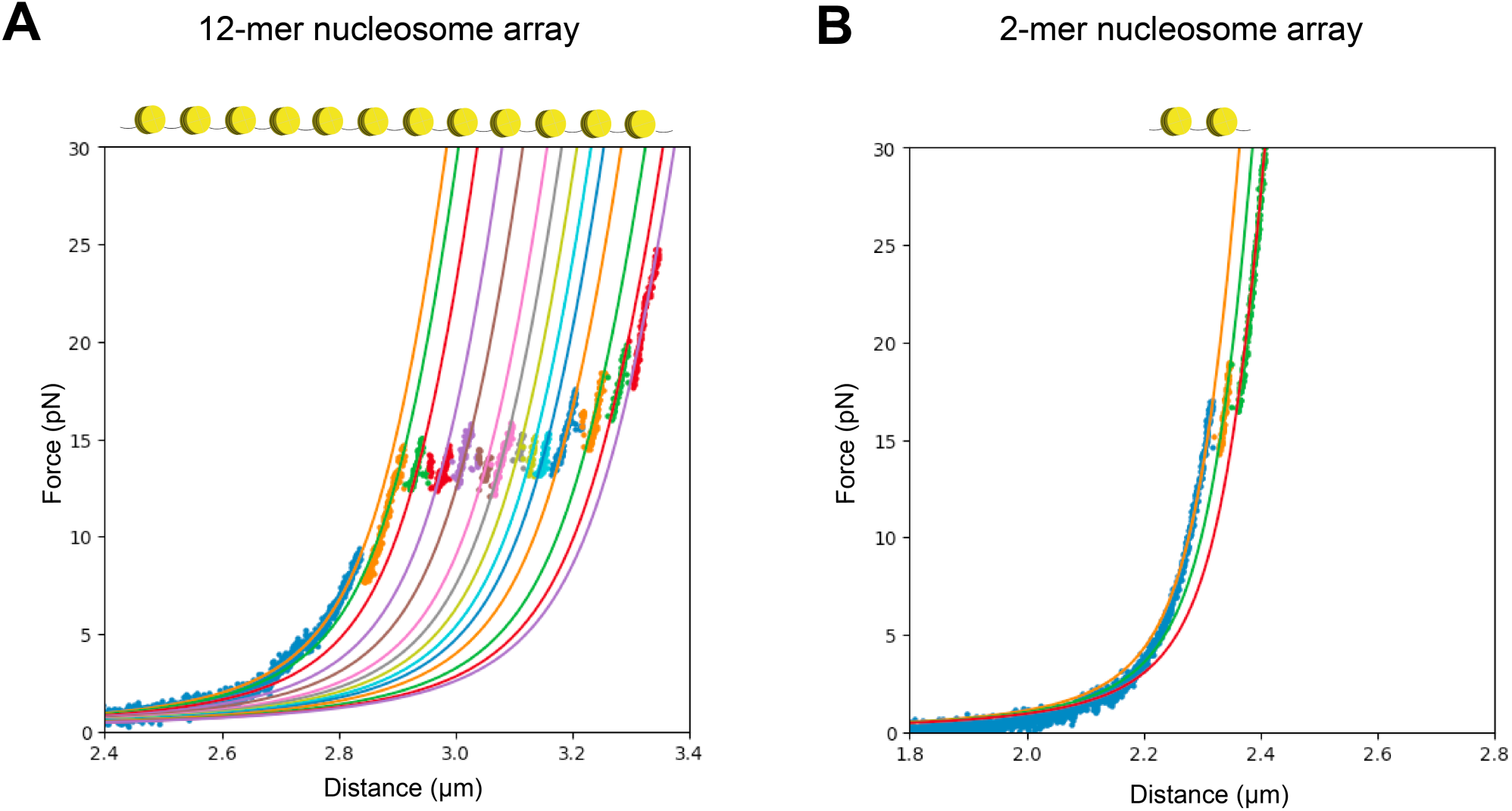
Segmentation analysis of force-extension curves. (**A**) A representative force-extension curve and its fitted segments for a 12-mer nucleosome array. (**B**) A representative force-extension curve and its fitted segments for a dinucleosome substrate. See *Materials and Methods* for fitting procedures.

**Figure S3.**
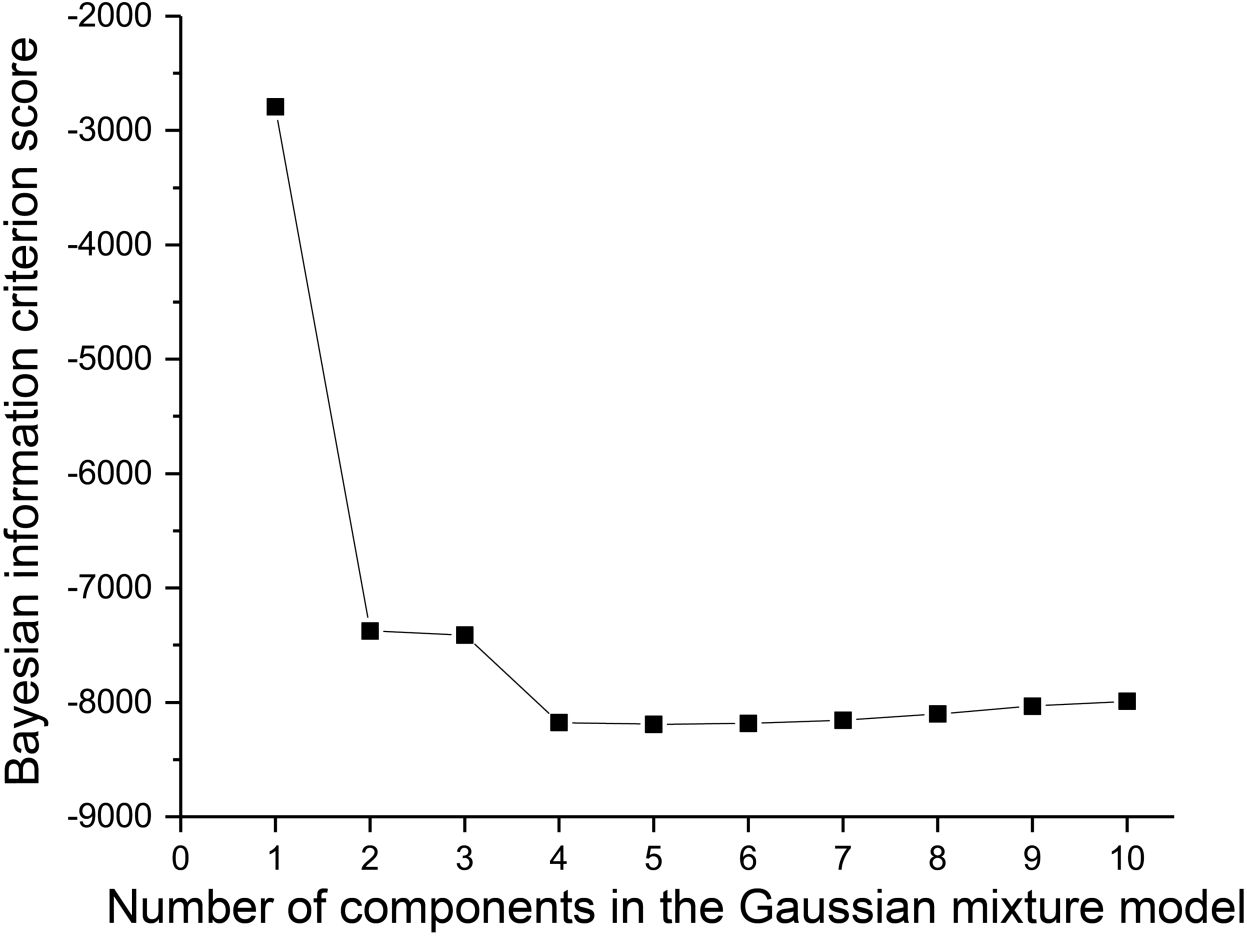
Cluster assignment for individual transitions in the force-extension curve. Bayesian information criterion (BIC) scores of Gaussian mixture models containing *N* components (*N* ranges from 1 to 10) to fit pooled data of Δ*L*_0_ (change in contour length per transition) derived from the force-extension curves. A four-component model was selected for all cluster assignments in this study because the BIC score ceased to improve when more components were added to the model. See *Materials and Methods* for details.

**Figure S4.**
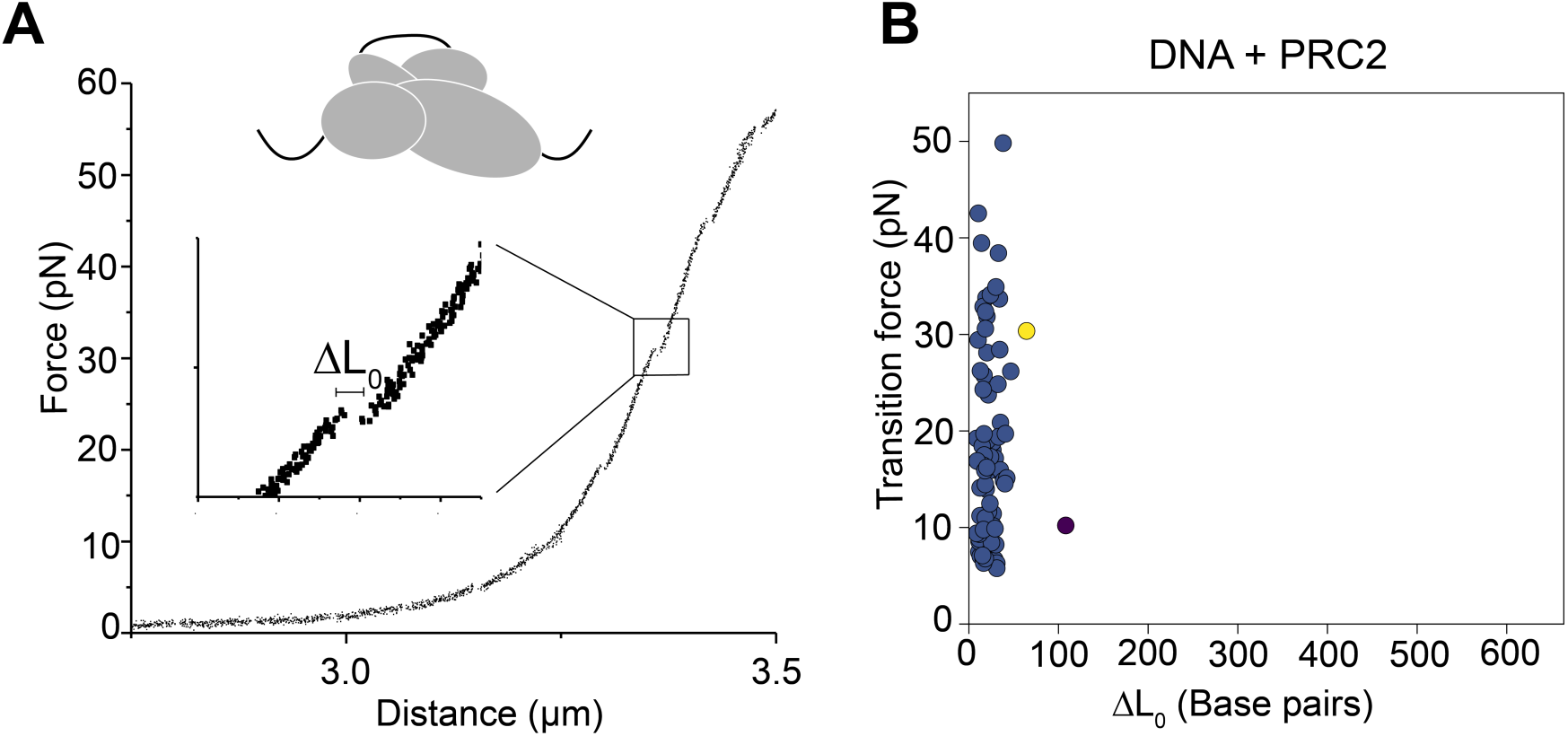
Analysis of PRC2–DNA interactions. (**A**) A representative force-extension curve of PRC2-bound bare DNA substrate and the zoomed-in view of one transition signifying PRC2 disengagement. (**B**) Cluster analysis of all transitions observed in the force-extension curves of PRC2-bound bare DNA.

**Figure S5.**
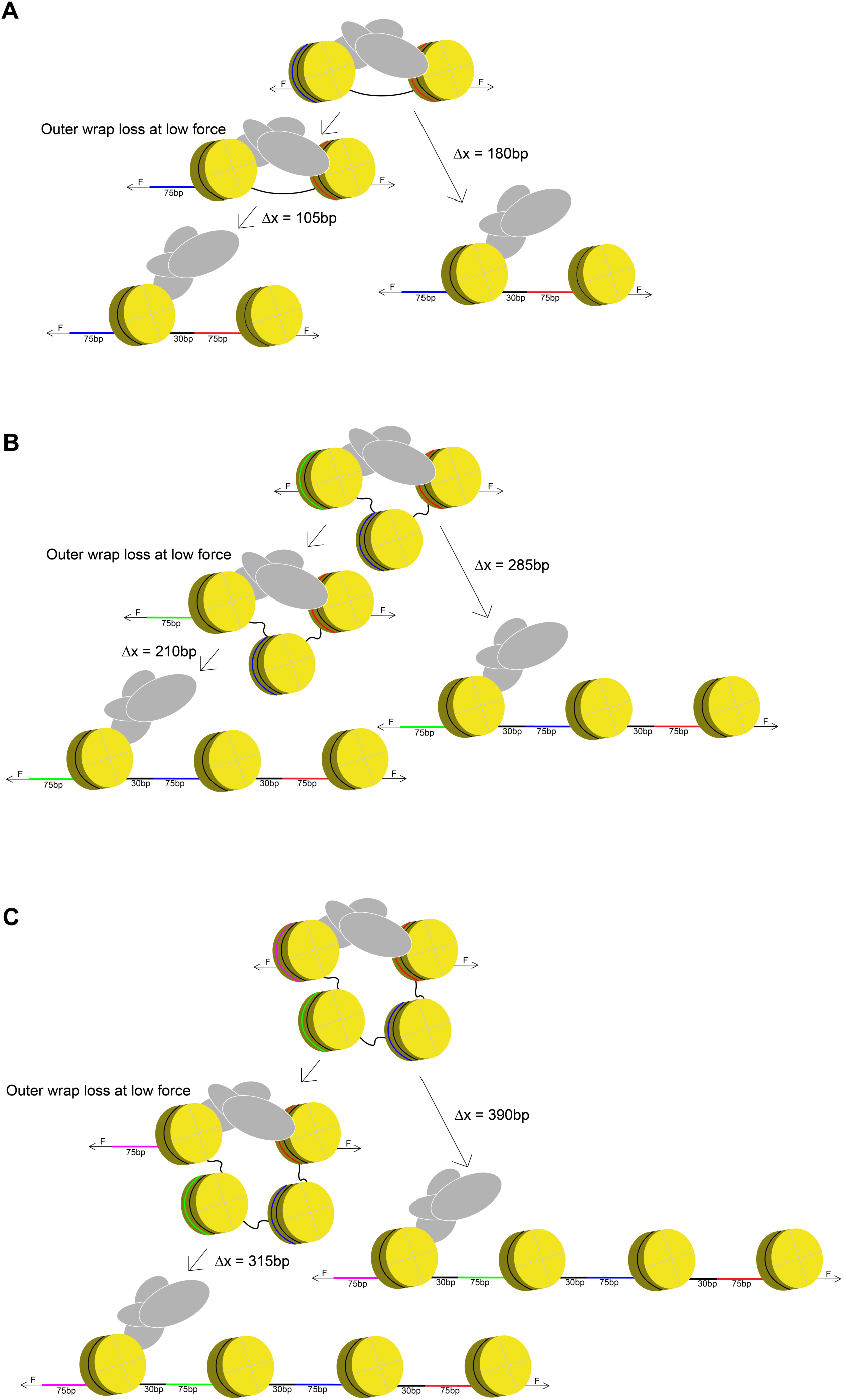
Scenarios for the amount of DNA released after disruption of PRC2-mediated bridging of nucleosome pairs. (**A**) Scenarios for force-induced disengagement of PRC2 that was initially engaging with two adjacent nucleosomes (Nucl_1-2_ mode). In the left pathway, the outer wrap of the first nucleosome (blue segment) is not sequestered by PRC2, thus undone at low forces. Upon PRC2 unbinding that occurs at a higher force, the outer wrap of the second nucleosome (red segment, ∼75 bp) and the 30-bp linker DNA—with a total length of ∼105 bp—are released. The inner wraps of the nucleosomes have similar stabilities to PRC2 engagement. Therefore they are expected to unravel later as independent transitions. Alternatively, as shown in the right pathway, PRC2 sequesters both outer wraps. In this scenario, PRC2 disengagement would release two outer wraps (∼150 bp) plus one linker DNA, totaling ∼180 bp. If PRC2 sequesters part of the first outer wrap, a number between 105 and 180 bp is expected for the amount of DNA released. (**B**) Same as (**A**), except for describing the binding mode in which PRC2 bridges two nucleosomes separated by one spacer nucleosome (Nucl_1-3_ mode). A total between 210 and 285 bp of DNA is expected to be released upon PRC2 disengagement. (**C**) Same as (**A**), except for describing the Nucl_1-4_ mode. A total between 315 and 390 bp of DNA is expected to be released upon PRC2 disengagement.

**Figure S6.**
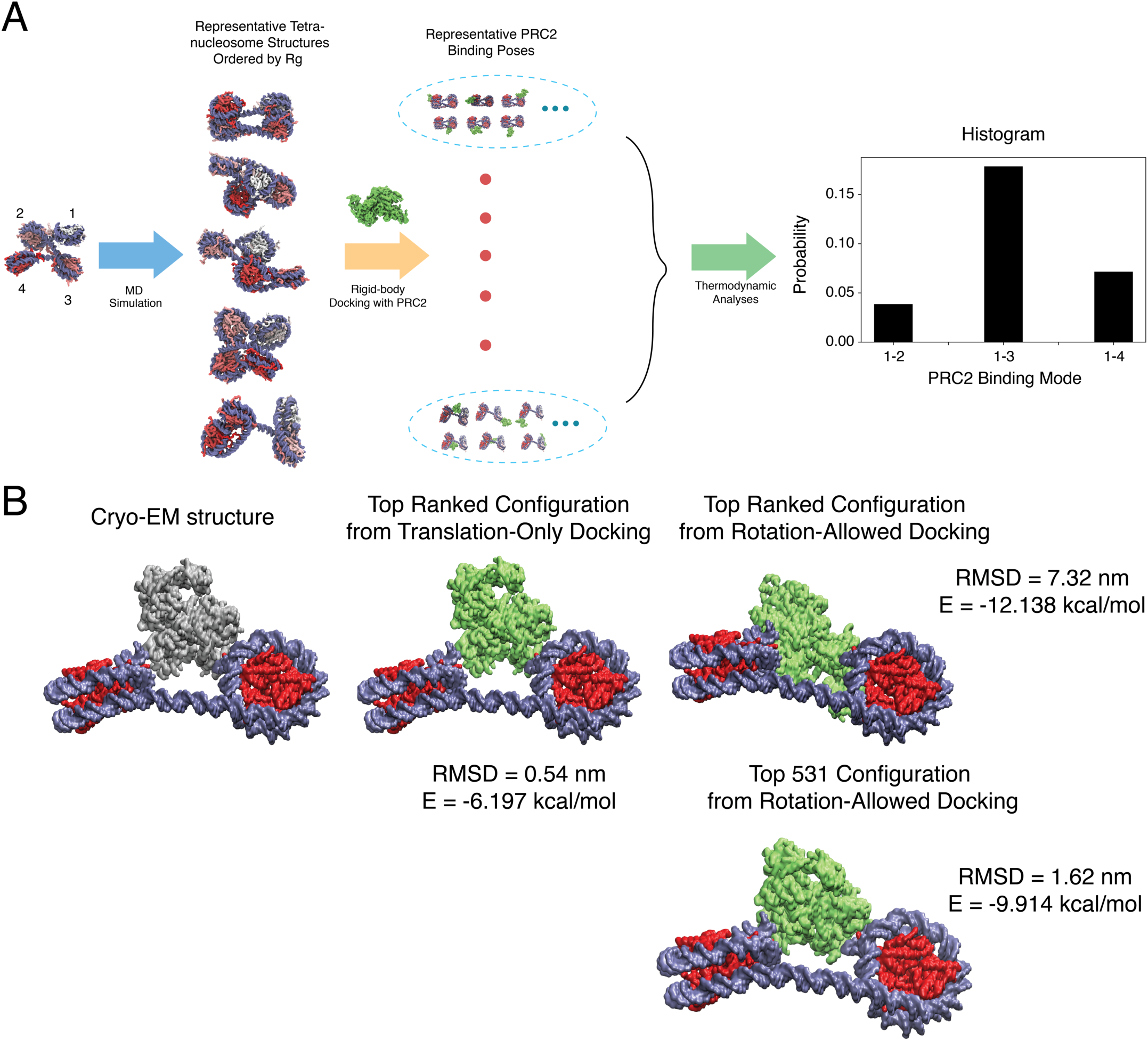
Coarse-grained modeling of PRC2–chromatin interactions. (**A**) Flowchart for the algorithm used to quantify the various binding modes PRC2 engaged with a tetranucleosome. See *Materials and Methods* for simulation details. (**B**) Comparison between experimental and simulated PRC2 binding poses. (*Left*) Cryo-EM structure of the PRC2-dinucleosome complex determined by Poepsel *et al.* (*6*). (*Middle*) Lowest energy configuration predicted from rigid docking using the coarse-grained force field detailed in *Materials and Methods*. The PRC2 structure used for docking includes SUZ12, EZH2, EED, and RBBP4 residues resolved in the cryo-EM structure. Only translational motion was allowed in these docking simulations and PRC2 was fixed in the same orientation as that found in the cryo-EM structure. The Cα RMSD between docked and cryo-EM structures is 0.54 nm. (*Right*) Lowest energy (top) and a low RMSD (bottom) configurations from docking simulations that allow both translational and rotational motions. In the lowest energy configuration, PRC2 is again juxtaposed between the two nucleosomes as found in cryo-EM, but with a slightly tilted orientation. The difference in orientation between simulated and experimental structures could be due to the omission of disordered regions in both PRC2 and histones during docking. The accuracy of force field could also potentially impact PRC2 orientation. In particular, we found that the top 531 ranked structure, though with an energy difference of less than 3 kcal/mol, is in much better agreement with the cryo-EM structure with a Cα RMSD of 1.62 nm. These results with the dinucleosome, therefore, support the use of the coarse-grained model for studying PRC2–chromatin binding modes.

**Figure S7.**
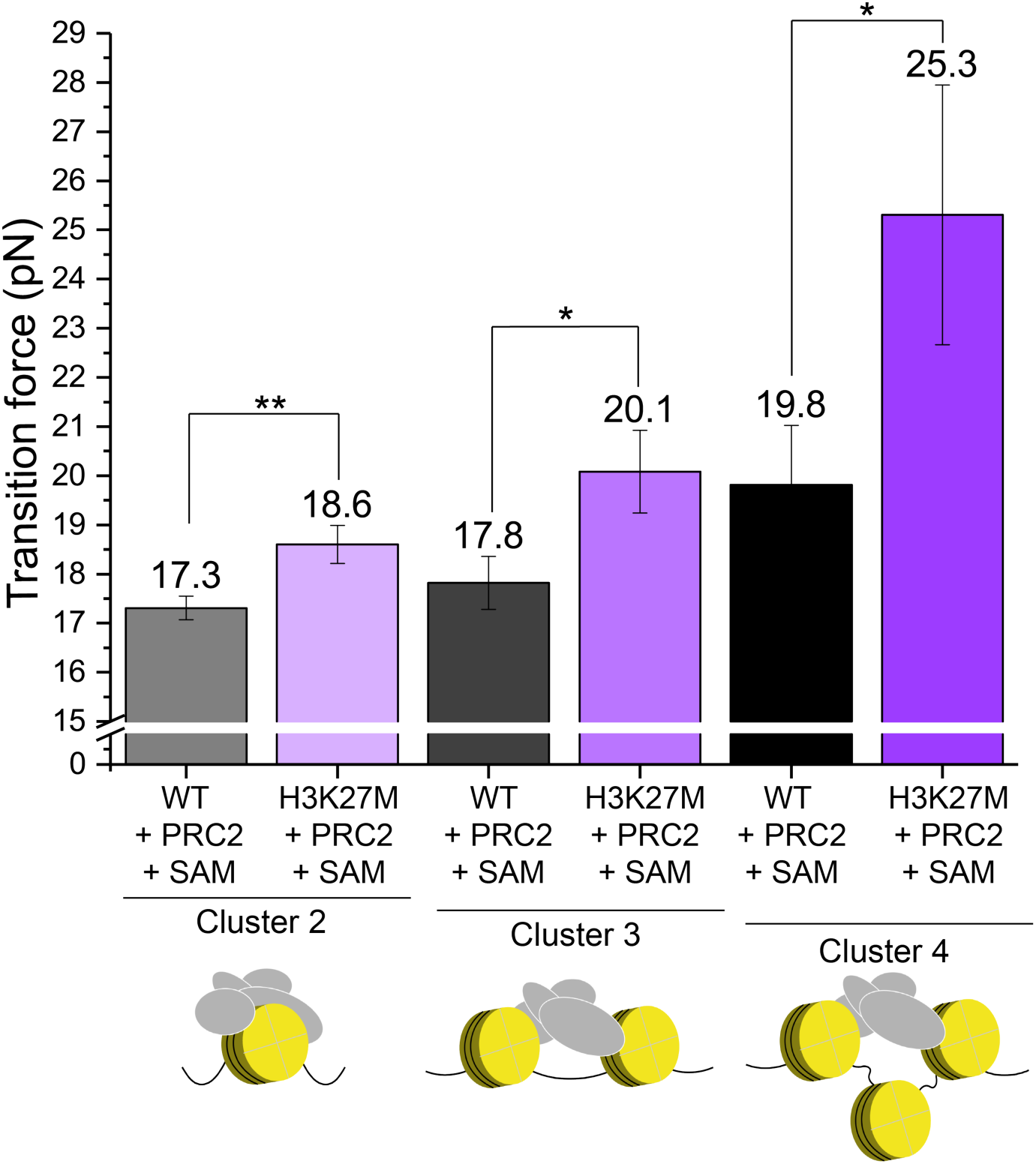
H3K27M mutation stabilizes PRC2–nucleosome interactions. In each cluster that corresponds to mononucleosome binding (Cluster 2), adjacent nucleosome bridging (Cluster 3), and non-adjacent nucleosome bridging (Cluster 4), a significantly higher average transition force was measured with 100% H3K27M-containing 12-mer arrays compared to wild-type arrays. Measurements were taken in the presence of SAM.

**Figure S8.**
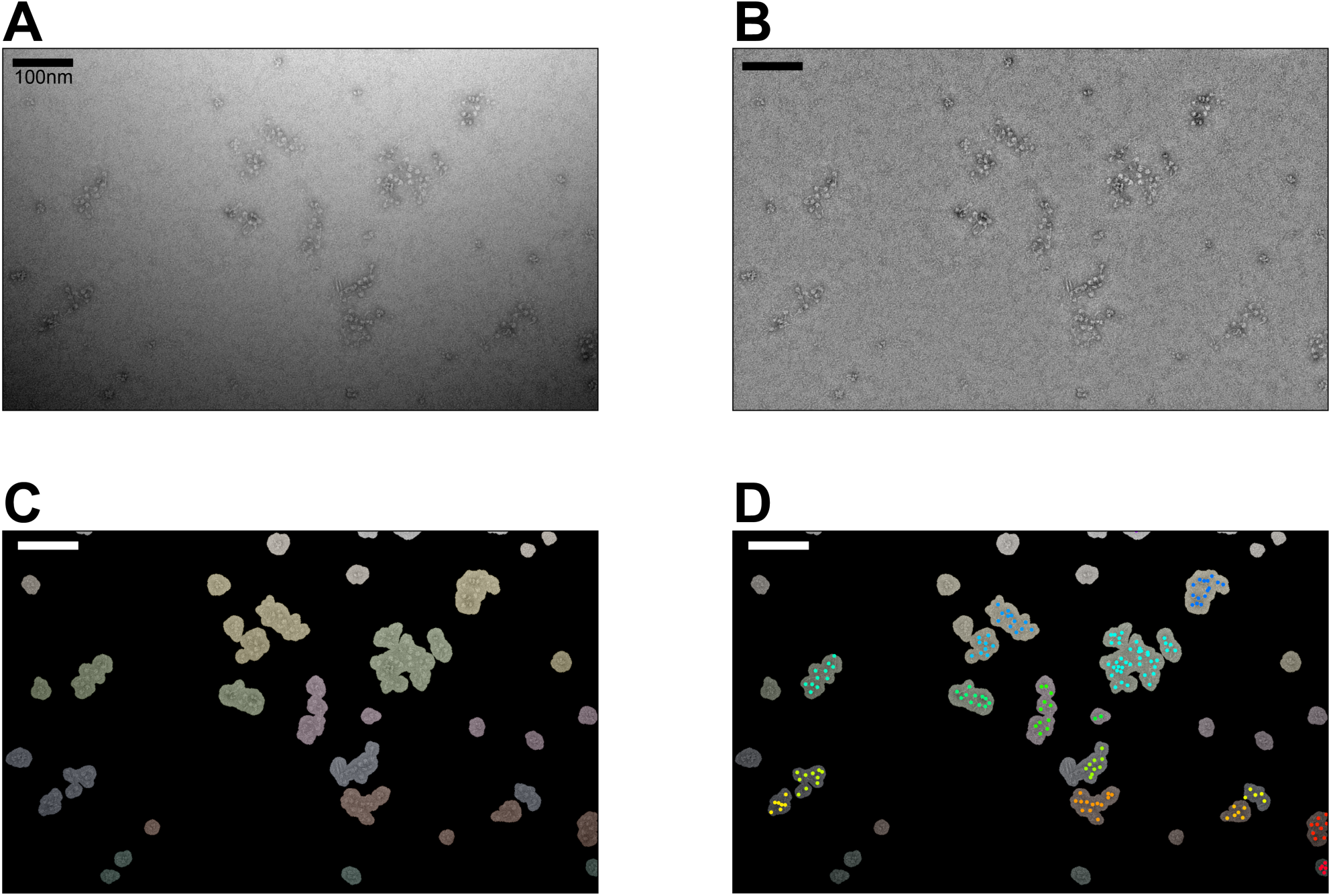
Workflow for analyzing the electron micrographs of nucleosome arrays. (**A**) A representative negative-stain EM micrograph of 12-mer nucleosome arrays mixed with PRC2 at a 1:1 ratio (150 nM each). (**B**) The same micrograph after illumination correction to achieve a uniform background. (**C**) Binary masks of individualized nucleosome arrays. (**D**) Manually picked nucleosomes (colored dots) inside their corresponding masks. Only masks associated with five or more picked nucleosomes were selected for further analysis to evaluate the level of compaction. See *Materials and Methods* for details.

